# Most High Throughput Expression Data Sets Are Underpowered

**DOI:** 10.1101/2022.08.03.502688

**Authors:** Alexander J. Trostle, Jiasheng Wang, Lucian Li, Ying-Wooi Wan, Zhandong Liu

## Abstract

Researchers handicap their high-throughput sequencing experiments when they do not perform enough biological replicates. Given that biological tissues can be highly variable, too few replicates will lead to false negatives, false positives, irreproducibility, and failure to detect real biological signatures. We propose that at least six biological replicates per condition becomes the standard for high-throughput expression datasets.

## Main

High-throughput sequencing is pervasive, routine, and cost effective. Signal-to-noise ratio should be considered in all quantitative experiments, but high-throughput sequencing is particularly vulnerable to high levels of background noise from non-biological sources. It is challenging to separate this technical noise from true biological variance. We will never completely eliminate all technical noise, but by increasing statistical power, we can improve the signal to noise ratio (SNR) of our data.

To examine signal and noise in biological data, we chose three experimental model organism datasets to ensure large total sample count and diversity of both platform and species: a human bulk transcriptome data set from distinct naïve pluripotent states, a nuclear bulk transcriptomic data set from a *MECP2* knockout mouse model, and a proteomics data set from a *Drosophila* model of Parkinson’s disease (based on alpha-synuclein mutation). These three datasets also represent different effect sizes: iPSC states show dramatically different gene expression profiles, *MECP2* knockout data has low gene expression effect size but perturbations are broad and subtle, and Alpha synuclein regulates cellular function via several different mechanisms, including protein expression [3]. These data sets have nine, ten, and eight control and experimental samples each, respectively, making them exemplary in their large *n* (sample size). By contrast, the median number of samples in the massive public database recount3 (2) is only four. Our selected large datasets also enable us to simulate under-sampled (low *n*) experiments and to quantify differences in the results. Population level data sets with huge *n* do exist, but for testing the null hypothesis that two groups of samples are not different in any way, these data sets are inappropriate [4].

Our analyses identified several specific limitations inherent to under-sampled high-throughput data. Lower sample number experiments yield fewer differentially expressed genes (DEGs) (**Figure 1, column 1**). Three-sample experiments can have orders of magnitude fewer statistically significant genes than the full data set at a 1% FDR threshold, and even just increasing the sample size from three to four can yield hundreds of additional genes. To better understand where saturation on DEG number versus *n* occurs, we repeated this analysis with very strict fold-change cutoffs (**Figure 1, column 2**). The human DEG number flattens out, but low sample numbers show high run-to-run variance, suggesting false positives. The mouse and fly data show a plateau effect at around six samples, where additional samples yield more DEGs but with diminishing returns. Power curves are plotted for DEGs in 30% or more repetitions of the down-sampled experiments (**Figure 1 column 3**). These show the full scope of the relation between DEG number, fold change, and sample size. Under-sampled datasets do not capture many low-fold change DEGs.

**Figure 1.**
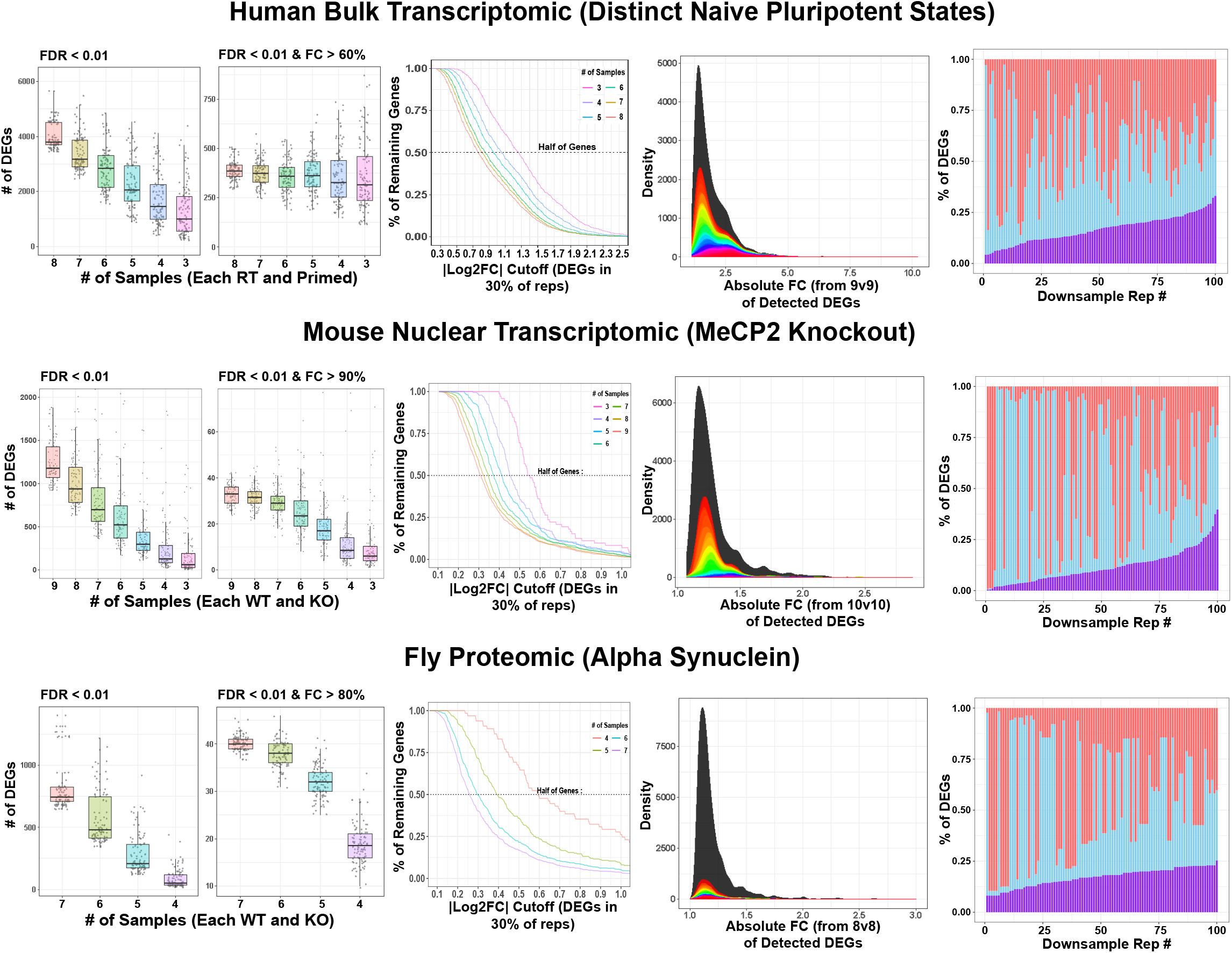
Down-sampling analysis across expression data sets. Each row is a data set. Column 1 shows the effect that *n* number has on DEG yield. Column 2 shows the same plots at a strict fold change cutoff. The power curve saturates and flattens out. Column 3 shows this continuous relationship between power and effect size with one line per down-sample. Column 4 shows effect size density curves for many low power down-samples in color and the “ground truth” in black. Column 5 is a visualization of the precision in low power data sets. With down-sampling, we simulate 100 paired low power experiments and plot the overlap of their results.

If we can still detect some DEGs even in under-sampled data, one might hope that these are the genes with the greatest changes in expression. To assess possible biases in which DEGs are not detected, **Figure 1 column 4** shows a black curve for the distribution of fold-change in the full experiment. This black density distribution is our best approximation to ground truth. Colored curves are then plotted on top for each of the 100 *n* = 3 down-samples. The distribution of the colored curves is similar to the full experiments in these datasets, suggesting that there is no clear bias in terms of fold-change. Unfortunately, this means that under-sampled data sets cannot reliably generate any particular subset of ground truth DEGs. If these lost DEGs were all low-fold change, researchers could at least interpret their low sample number results with that limitation in mind. The repetitions also differed significantly from one another, further demonstrating the high variance of an under-sampled experiment.

Reproducibility has become a concern across scientific fields, as many highly cited studies are not replicable [1]. High variance in DEG number from under-sampled data suggests that these few detected DEGs would not be reproducible. To examine this possibility, we simulated independent datasets of identical experimental design. Out of the full sample space, we generated 200 new down-samplings of *n* = 3 with paired non-overlapping samples. This methodology splits the ten samples up into distinct 3v3 repetitions, as if two separate labs each generated an under-sampled data set and are now interested in comparing them. We analyzed the repetitions and plotted the DEG overlap between matched pairs (**Figure 1, column 5**). Between a third and a quarter of genes were recovered as overlaps. This simulation is a best-case scenario; researchers looking to validate or compare under-sampled data sets should not expect as much DEG congruency. The 25-33% overlapping DEGs are almost all present in the full sample space DEG list, but unfortunately, this overlap constitutes less than 5% of the “ground truth” DEGs. Researchers with access to several low-power datasets can have some trust in the few overlapping DEGs they observe, but individual under-sampled experiments will yield false positives.

To get a broad empirical sense of the relationship between statistical power, *n* number, effect size, and biological variance, we have also down-sampled 461 expression data sets from recount3. We use this unbiased, large scale power analysis to develop a more objective recommendation for researchers. On curves of median DEG number per down-sample, we calculate where the second derivative is equal to 0. This should generally represent a pivot point and indicate where saturation on power begins. We calculate this across effect sizes and plot the density (**Figure 2**). Results clearly show that *n*=6 is where saturation begins. Choosing 6 is worthwhile because every extra sample between 3 and 6 has a sizable improvement on the conclusions of the experiment. Researchers with spare funding will benefit from generating even more biological replicates. Power analysis on a *Saccharomyces cerevisiae* Δ*snf2* model by Schurch et al. (2016) corroborates our findings.

**Figure 2.**
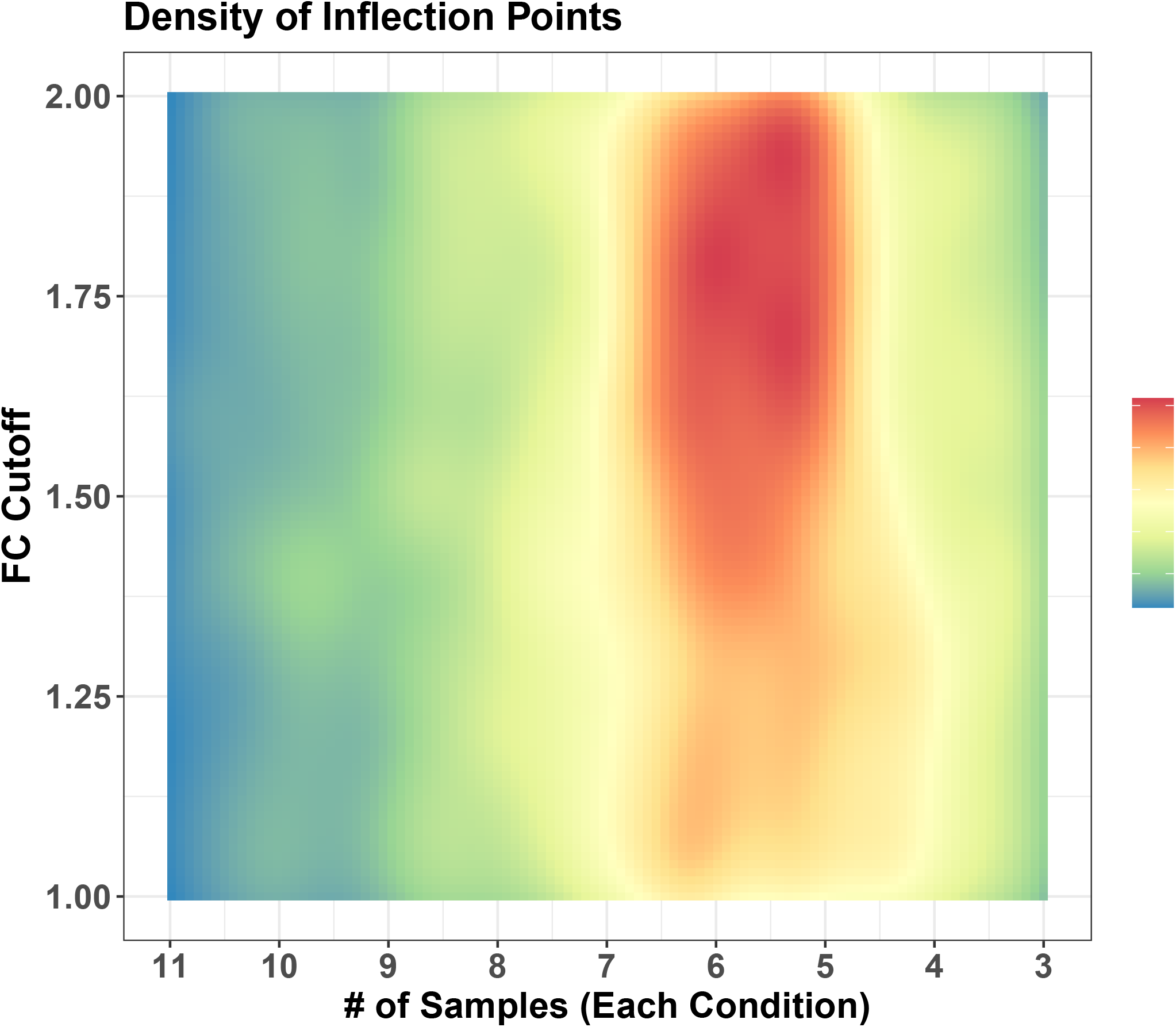
Density of power saturation across many expression data sets. Saturation / pivot point is shown across down-samples and FC thresholds.

These findings are robust across platform, effect size, and species. Six or more samples per condition needs to become the new standard for publication-quality datasets. Researchers generating high throughput data need more samples to avoid fragmentary, highly variable, and unreliable results. The quantitative advantages of just a few extra samples are clear.

## Methods

All analysis was conducted in R version R version 3.5.2 (Eggshell Igloo). Plots were made with ggplot2 R package version 3.2.1. Mouse samples were aligned to GENCODE GRCm38p6 primary assembly, version 18 (https://www.gencodegenes.org/mouse/release_M18.html), with STAR version 2.6.0a at default parameters. The assembly also contained an appended copy of human MeCP2 from hg38. We assessed alignment quality with RSeQC geneBody_coverage and read_distribution, both version 3.0.0. DEG analysis was performed in with DESeq2 version 1.24.0 [4], after loose expression filtering (per data set, a gene must have a sum of 10 counts in at least half the samples). Analysis by Schurch et al. (2016) recommends DESeq2. Human samples were from studies treated by recount3 pipeline [2]. They used the UCSC hg38 assembly based on GRCh38, with STAR version 2.73a. For QC and controls, many tools have been used including seqtk, the idxtats subcommand of samtools, the output of STAR, Megadepth tool, and featureCounts (http://rna.recount.bio/docs/quality-check-fields.html). The DEG analysis were also performed in with DESeq2 version 1.24.0, after loose expression filtering (per data set, a gene must have a sum of 20 counts in at least half the samples). Fly samples were generated and processed as described in Yu et. Al (2022).

The inflection point analysis was conducted by downloading RNAseq data sets from recount3. Recount data was filtered on *n* number between 8 and 12 and then on full data DEG yield between 300 and 6000. This results in 461 data sets. We down-sample them as detailed above, and calculate where the second derivative of their median DEG number curves equals 0 with R package *akmedoids* version 1.3.0. Calculation is done for FC cutoffs in 10% intervals between 1 and 2. When multiple 0s of the second derivative were found, the one at lower *n* number was chosen. Inflection points were successfully called for 450 data sets and then plotted in figure 2 using stat_density_2d from R package *ggplot2* version 3.3.3.

## Acknowledgments

We would like to thank Dr. Hari Krishna Yalamanchili for helpful comments and insights.

## Notes

### Competing Interest Statement

The authors have declared no competing interest.

